# Kirrel3-mediated synapse formation is attenuated by disease-associated missense variants

**DOI:** 10.1101/2019.12.30.891085

**Authors:** Matthew R. Taylor, E. Anne Martin, Brooke Sinnen, Rajdeep Trilokekar, Emmanuelle Ranza, Stylianos E. Antonarakis, Megan E. Williams

## Abstract

Missense variants in Kirrel3 are repeatedly identified as risk factors for autism spectrum disorder and intellectual disability but it has not been reported if or how these variants disrupt Kirrel3 function. Previously, we studied Kirrel3 loss-of-function using knockout mice and showed that Kirrel3 is a synaptic adhesion molecule necessary to form one specific type of hippocampal synapse in vivo. Here, we developed a new gain-of-function assay for Kirrel3 and find that wild-type Kirrel3 induces synapse formation selectively between Kirrel3-expressing cells via homophilic, trans-cellular binding. We tested six disease-associated Kirrel3 missense variants and find that five attenuate this synaptogenic function. All variants tested traffic to the cell surface and localize to synapses similar to wild-type Kirrel3. Two tested variants lack homophilic trans-cellular binding, which likely accounts for their reduced synaptogenic function. Interestingly, we also identified variants that bind in trans but cannot induce synapses, indicating Kirrel3 trans-cellular binding is necessary but not sufficient for its synaptogenic function. Collectively, these results suggest Kirrel3 functions as a synaptogenic, cell-recognition molecule, and this function is attenuated by missense variants associated with autism spectrum disorder and intellectual disability. Thus, we provide critical insight to Kirrel3 function in typical brain development and the consequences of missense variants associated with autism spectrum disorder and intellectual disability.

**SIGNIFICANCE STATEMENT:** Here, we advance our understanding of mechanisms mediating target-specific synapse formation by providing evidence that Kirrel3 trans-cellular interactions mediate contact recognition and signaling to promote synapse development. Moreover, this is the first study to test the effects of disease-associated Kirrel3 missense variants on synapse formation, and thereby, provides a framework to understand the etiology of complex neurodevelopmental disorders arising from rare missense variants in synaptic genes.

## INTRODUCTION

Autism spectrum disorder (ASD) has a strong genetic basis but the mechanistic links between genetic risk variants and brain development remain poorly understood (Tick et al., 2016; Sandin et al., 2017; Bai et al., 2019). This is likely because ASD patients have a heterogeneous set of atypical behaviors presenting with a variety of co-morbid conditions and genetic variants in hundreds of genes are identified as ASD risk factors (American Psychiatric Association, 2013; Rubeis et al., 2014; Iossifov et al., 2014; Constantino and Charman, 2016). As such, it is important to understand how different types of genetic risk variants functionally impact cells, circuits, and behavior to properly understand, diagnose, and treat the diverse etiologies of individual patients (Geschwind and State, 2015).

Kirrel3 is an immunoglobulin (Ig)-domain containing transmembrane protein (Fig. 1A) that undergoes homophilic trans-cellular binding (Sellin et al., 2002; Gerke et al, 2005; Martin et al., 2015). Importantly, copy number variations and missense variants in Kirrel3 are repeatedly associated with neurodevelopmental disorders including ASD and intellectual disability (ID) (Table 1) (Bhalla et al, 2008; Kaminsky et al, 2011; Ben-David and Shifman, 2012; Guerin et al, 2012; Michaelson et al, 2012; Neale et al, 2012; Talkowski et al, 2012; Iossifov et al., 2014; Rubeis et al, 2014; Wang et al, 2016; Yuen et al, 2016; Li et al, 2017; Guo et al, 2019; Leblond, et al, 2019). There are currently at least 17 missense variants identified from multiple, independent studies lending credence to the possibility that altering Kirrel3 function leads to ASD and ID. However, it remains unknown if Kirrel3 missense variants affect protein function.

**Table 1:** Kirrel3 missense variants associated with neurodevelopmental disorders.

| Variant ID | Amino Acid Change | Associated Conditions | Reference |
| --- | --- | --- | --- |
| rs182260035 | Val30Met | ASD | Rubeis et al, 2014 |
| rs119462978 | Arg40Trp | ID | Bhalla et al, 2008 |
| rs779979146 | Arg161His | ASD | Iossifov et al, 2014; Yuen et al, 2016 |
| rs376743687 | His171Gln | ASD | Rubeis et al, 2014 |
| rs572991045 | Arg205Gln | ASD | Rubeis et al, 2014 |
|  | Lys212Asn | ASD | Rubeis et al, 2014 |
| rs372786849 | Ser241Leu | ASD | Li et al, 2017 |
| rs201725914 | Val303Ala | ASD | Rubeis et al, 2014 |
| rs114378922 | Arg336Gln | ID | Bhalla et al, 2008 |
|  | Trp358Cys | ASD | Li et al, 2017 |
| rs201882059 | Ala393Thr | ASD | Wang et al, 2016 |
| rs369363937 | Ala393Val | ASD | Wang et al, 2016 |
| rs1053637755 | Gln428Lys | ASD | Li et al, 2017 |
| rs760499655 | Pro446Leu | ASD | Li et al, 2017 |
|  | Arg562Leu | ASD | Leblond, et al, 2019 |
| rs773312950 | Arg650His | ASD | Guo et al, 2019 |
| rs1057519593 | M6731I | ID, ADHD | This study |
| rs119462980 | Val731Phe | ID | Bhalla et al, 2008 |

**Figure 1.**
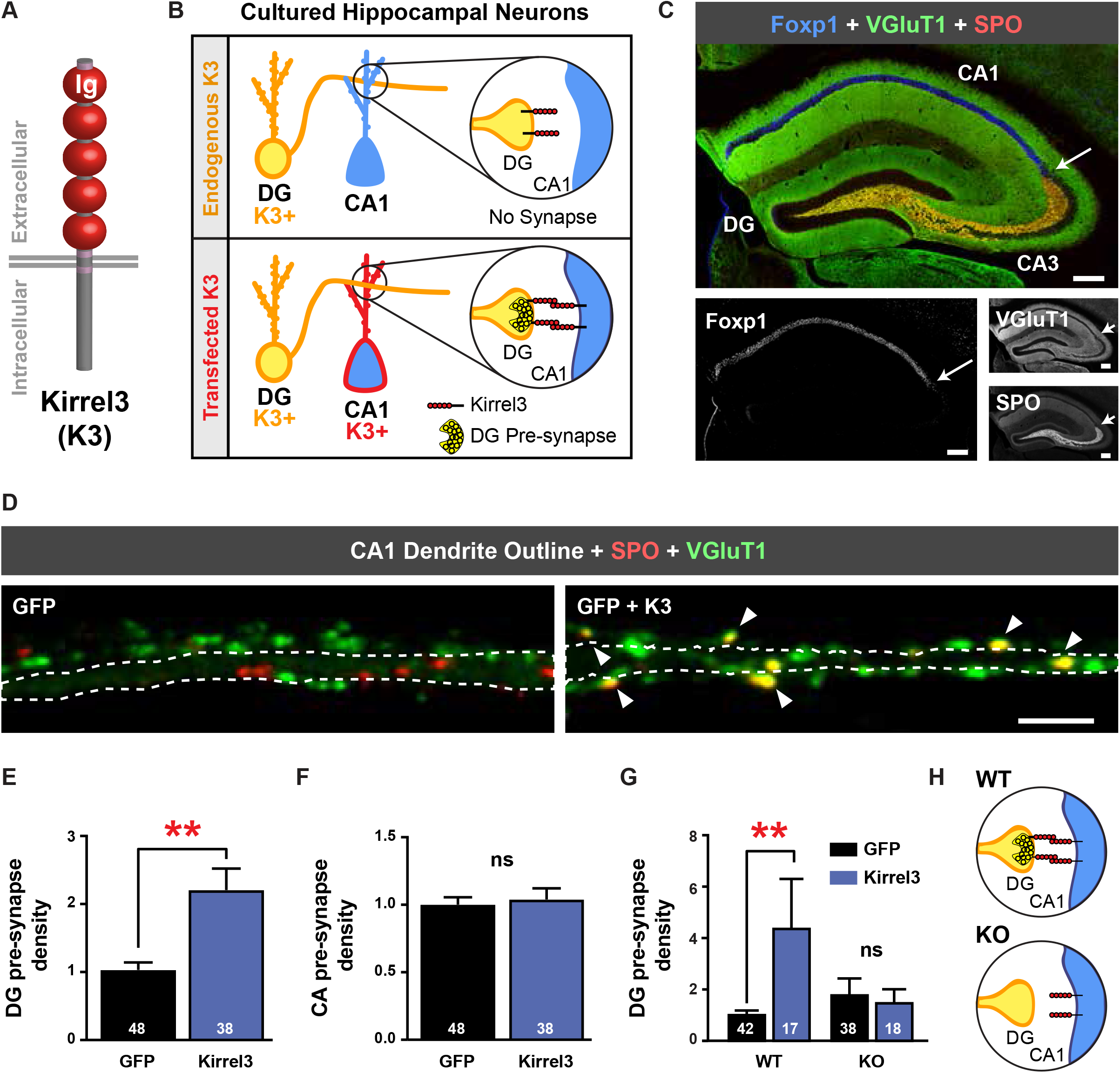
Ectopic Kirrel3 expression induces ectopic DG pre-synapses in cultured neurons. **A**, Schematic of Kirrel3 (K3) protein. Ig; Immunoglobulin domain. **B**, Schematic of the gain-of-function assay design. Top: In the control condition, a Kirrel3 positive DG neuron contacts a Kirrel3 negative CA1 dendrite and does not form a pre-synapse. Bottom: A Kirrel3 positive DG neuron contacts a Kirrel3-transfected CA1 neuron allowing for a DG pre-synapse to form. **C**, Tiled 20x image of an adult mouse hippocampal section immunostained with the CA1 marker Foxp1 (blue) and the presynaptic markers VGluT1 (green) and SPO (red). Grayscale images below. In each image the white arrow indicates approximate CA3 border. Scale bars = 200μm. **D**, Dendrites from cultured rat CA1 dendrites transfected with GFP (left) or GFP + Kirrel3 (right). GFP is not shown but is outlined in white. DG pre-synapses (yellow) are identified by co-labelling of SPO (red) and VGluT1 (green). Arrows point to DG pre-synapses. Scale bar = 5 μm. **E**, Quantification of DG pre-synapse density per length of CA1 dendrite and normalized to the GFP condition. n=38-48 neurons (indicated in each bar) from 3 cultures, p=0.0027 (Mann Whitney test). **F**, Quantification of CA synapse density per length of CA1 dendrite and normalized to the GFP condition. n=38-48 neurons (indicated in each bar) from 3 cultures, p=0.9552 (Mann Whitney test). **G**, Quantification of DG pre-synapse density per length of CA1 dendrite from Kirrel3 wild-type (WT) and knockout (KO) cultures. All are normalized to the wild-type GFP condition. n=17-42 neurons (indicated in each bar) from 3 cultures, WT: GFP vs GFP+Kirrel3 p=0.0088; KO: GFP vs GFP+Kirrel3, p=0.9560 (Two-way ANOVA with Sidak’s multiple comparisons, p= 0.0266; interaction) **H**, Schematic of WT versus KO assay results suggesting that Kirrel3 must be present in both the pre- and postsynaptic neuron to induce DG synapse formation. For F, and G, ns = not significant.

We recently discovered Kirrel3 is necessary for the formation of a specific type of synapse in the rodent hippocampus (Martin et al., 2015; Martin et al., 2017). Here, dentate granule (DG) neurons project axons, called mossy fibers, to area CA3 where they make unique excitatory synaptic complexes. These mossy fiber synapse complexes connect DG neurons to CA3 neurons via giant presynaptic boutons, each of which contains dozens of active zones capable of vesicle release (Acsady et al., 1998; Bischofberger, 2006). Each giant presynaptic bouton also projects several presynaptic filopodia that, in turn, build smaller excitatory synapses onto both CA3 neurons and GABAergic neurons in area CA3 (Acsády et al. 1998; Chicurel and Harris, 1992; Martin et al., 2017). In the hippocampus, Kirrel3 is only detected in DG neurons and GABAergic neurons (Martin et al., 2015). Consequently, Kirrel3 knockout animals have a highly specific loss of DG to GABAergic neuron mossy fiber filopodial synapses, presumably via loss of trans-cellular contact recognition and downstream signaling mediated by Kirrel3 homophilic binding (Martin et al., 2015; Martin et al., 2017). Ultimately Kirrel3 loss-of-function leads to a loss of feed-forward inhibition and an excitation/inhibition imbalance in the developing hippocampus (Martin et al., 2015). In sum, previous work provides strong evidence that Kirrel3 is an important target-specific, pro-synaptogenic molecule in vivo.

Several other recent studies also support a general role for Kirrel3 in mammalian neurodevelopment. Specifically, Kirrel3 may also function at post-synapses in DG dendrites as DG neurons from Kirrel3 knockout mice have more frequent miniature excitatory postsynaptic events in young animals (Roh et al., 2017). Furthermore, Kirrel3 knockout mice have an array of aberrant behavioral phenotypes, most notably characterized by hyperactivity, social, and communication deficits (Choi et al., 2015; Hisaoka et al., 2018; Völker et al., 2018). Taken together, it is becoming clear that lack of Kirrel3 function alters synapse development and adversely affects circuit function and behavior, but the mechanisms underlying Kirrel3 function remain unknown.

Here, we sought to advance our understanding of Kirrel3 function and investigate how Kirrel3 variants could cause disease. To accomplish this, we developed a gain-of-function in vitro assay and discovered that ectopic Kirrel3 expression specifically induces ectopic DG pre-synapse formation via Kirrel3 homophilic interactions. We then tested the function of several disease-associated Kirrel3 missense variants identified in humans using the gain-of-function assay. We found that five out of six disease-associated Kirrel3 variants tested have attenuated synaptogenic function. Intriguingly, we identified two classes of non-functional variants: one that can neither mediate homophilic trans-cellular interactions nor induce synapses, and one that can mediate homophilic trans-cellular interactions yet has impaired synaptogenesis. We conclude that trans-cellular binding is necessary but not sufficient for Kirrel3-mediated synapse formation and provide strong support that most disease-associated Kirrel3 variants impair Kirrel3 function and are bona fide risk factors for neurodevelopmental disorders.

## MATERIALS and METHODS

### Kirrel3 variant identification and cloning

Single nucleotide polymorphism (SNP) accession numbers for human disease-associated Kirrel3 missense variants are shown in Table 1 and were taken from the literature as indicated in the table and the SFARI gene database (https://gene.sfari.org; Banerjee-Basu and Packer, 2010; Abrahams et al, 2013). The M673I variant is newly discovered and described below. Human Kirrel3 isoform 1 (NM_032531.4 nucleotide; NP_115920.1 protein) and mouse Kirrel3 Isoform A (NM_001190911.1 nucleotide; NP_001177840.1 protein) were aligned using Clustal Omega (Madeira et al., 2019) and found to share 97.94% amino acid identity with identical length. Thus, all variants are named and numbered according these isoforms. However, all experiments in this study use a cDNA encoding mouse Kirrel3 isoform B (NM_026324.3 nucleotide; NP_080600.1) that is identical to isoform A except that it lacks an alternatively spliced micro-exon encoding 12 amino acids near the transmembrane domain. Isoform B contains all the conserved missense variant sites and was previously cloned by our lab with an extracellular FLAG-tag in the vector pBOS (Martin et al., 2015). Variants were cloned using Q5 Site Directed Mutagenesis (NEB Cat# E0554S) and QuikChangeII XL Site-Directed Mutagenesis (Agilent Technologies Cat# 200521-5) kits. Mouse cDNA codon changes were made as follows: R40W (CGG to TGG), R161H (CGG to CAT), R205Q (CGA to CAA), R336Q (CGA to CAA), M673I (ATG to ATT), and V731F (GTC to TTC).

### Discovery of the Kirrel3 M673I variant

We evaluated a 12 years old boy for ID, ADHD, and obesity. He is the only child of healthy unrelated parents, born at term after an uneventful pregnancy; growth parameters at birth were normal (height −0.5 SD, weight −0.5 SD and head circumference 0.5 SD). Neonatal hypotonia was noted. The patient subsequently showed global developmental delay, with walking at 26 months and marked language delay (first words at 4 years old). Neuropsychological evaluation at age 12 years showed moderate intellectual disability. He is integrated in specialized schooling and cannot count or read. He has fine motricity deficits. Moreover, he suffers from marked ADHD and benefits from methylphenidate treatment. There are no autistic traits. Since 8 years of age, he has behavioral problems with frequent oppositional crises. In addition, the parents report hyperphagia and food craving. At 12 years of age, his height is at 0.5 SD, for a weight of + 3SD (BMI 25, with truncal obesity) and head circumference at −1.5 SD. The patient also presents with mild dysmorphic features, i.e. exophthalmos, bulbous nose with slightly anteverted nares, thin upper lip and wide spaced teeth with two missing superior canines. Hands are normal except for the presence of fetal pads; feet show 2-3rd toe syndactyly. There are no transit or sleep issues. The boy had frequent otitis media during infancy, needing trans-tympanic drains; he is wearing glasses for mild hypermetropia and astigmatism. Cerebral MRI at age 2 showed mild temporal cortical atrophy and wide asymmetric aspect of posterior ventricular horns. Metabolic screening was normal as well as karyotype, FMR1 gene analysis, 15q11.2 DNA methylation, and array-CGH.

Exome sequencing followed by targeted analysis of a panel of 990 genes implicated in intellectual disability, identified a novel de novo missense variant in KIRREL3 c.2019G>A:p.(M673I) which was classified as likely pathogenic according to the ACMG guidelines (Richards et al, 2015). This variant has not been observed in gnomad V2.1.1 (https://gnomad.broadinstitute.org/gene/ENSG00000149571?dataset=gnomad_r2_1; 248436 alleles), or gnomAD V3 (https://gnomad.broadinstitute.org/gene/ENSG00000149571?dataset=gnomad_r3; 143348 alleles), or the Bravo databases (https://bravo.sph.umich.edu/freeze5/hg38/gene/ENSG00000149571; 125568 alleles) but a similar variant, M673V, has been observed once in gnomAD V2.1.1, twice in gnomAD V3, and twice in Bravo.

### Animals

Experiments involving animals were conducted in accordance with National Institute of Health guidelines with approval from the University of Utah Institutional Animal Care and Use Committee. Timed pregnant Sprague Dawley dams were ordered from Charles River (Strain Code 400; RRID: RGD_737926). Wild-type C57BL/6J mice (RRID: IMSR_JAX:000664) were originally obtained from The Jackson Laboratory. Generation of Kirrel3 knockout mice was previously reported (Prince et al., 2013). Knockouts were maintained in a C57BL/6J background. Hippocampi from male and female newborn (P0) pups were combined for neuron cultures.

### Immunostaining

Cells on glass coverslips were fixed at room temperature for 10-15 minutes in 4% PFA made in PBS. Cells were rinsed two times with PBS and blocked/permeabilized in PBS with 3% BSA and 0.1% Triton-x for 30 minutes. Cells were incubated with primary antibodies diluted in PBS with 3% BSA for 1-2 hours at room temperature with rocking, rinsed three times in PBS with 3% BSA and incubated with secondary antibodies diluted in PBS with 3% BSA for 45 minutes. After rinsing, coverslips were mounted in Fluoromount-G (Southern Biotech Cat# 0100-01). For surface labelling, live neurons were incubated in serum-free media at 37°C in a 5% CO_2_ incubator with chicken-anti-FLAG for 20 minutes. After initial incubation, neurons were fixed, permeabilized and stained as above.

### Antibodies

Primary antibodies used include: rabbit-anti-Fox1 (1:3600, Abcam, Cat# ab16645, RRID: AB_732428), guinea pig-anti-VGluT1 (1:5000, Millipore, Cat# AB5905, RRID: AB_2301751), goat-anti-synaptoporin (SPO, 1:1000, Santa Cruz Biotechnology, Cat# sc-51212, RRID: AB_ 1129844), goat-anti-GFP (1:3000, Abcam, Cat# ab6673, RRID: AB_305643), mouse-anti-PSD-95 (1:1000, Neuromab, Cat# 75-348, RRID: AB_2315909), chicken-anti-FLAG (1:1000, Gallus Immunotech, Cat# AFLAG), mouse-anti-FLAG M2 (1:5000, Sigma, Cat# F1804, RRID: AB_262044), mouse anti-FLAG M2-Cy3 conjugated (1:1000, Sigma, Cat# A9594, RRID: AB_ 439700), rabbit-anti-2A (1:1000, Millipore, Cat# ABS31, RRID: AB_11214282).

Secondary antibodies were all all used at 1:1000 and include: donkey-anti-rabbit-Alexa 488 (Invitrogen, Cat# A21206, RRID: AB141708), donkey-anti-rabbit-Cy3 (Jackson ImmunoResearch, Cat# 711-165-152, RRID: AB_2307443), donkey-anti-guinea pig-Alexa 647 (Jackson ImmunoResearch, Cat# 705-605-147, RRID: AB_2340437), donkey-anti-guinea pig-DyLight 405 (Jackson ImmunoResearch, Cat# 706-475-148, RRID: AB_2340470), donkey-anti-goat-Cy3 (Jackson ImmunoResearch, Cat# 705-165-147, RRID: AB_2307351), donkey-anti-goat-Alexa 647 (Jackson ImmunoResearch, Cat# 705-605-147, RRID: AB_2340437), donkey-anti-goat-Alexa 488 (Jackson ImmunoResearch, Cat# 705-545-147, RRID: AB_2336933), donkey-anti-mouse-Cy3 (Jackson ImmunoResearch, Cat# 715-165-150, RRID:AB_2340813), donkey-anti-mouse-Alexa 647 (Jackson ImmunoResearch, Cat# 715-605-150, RRID: AB_2340862), donkey-anti-mouse-Alexa 488 (Invitrogen, Cat# A21202, RRID: AB_141607), donkey-anti-chicken-Alexa 647 (Jackson ImmunoResearch, Cat# 703-605-155, RRID: AB_2340379), donkey-anti-chicken-Cy3 (Jackson ImmunoResearch, Cat# 703-165-155, RRID: AB_2340363).

### Neuron culture

Neuron cultures were prepared as previously described (Martin et al., 2015; Basu et al., 2017). Briefly, rat cortical astrocytes were plated to glass coverslips coated with 0.03mg/ml (~7.5μg/cm^2^) of PureCol (Advanced Biomatrix, Cat# 5005) and grown to confluence in glia feeding media: MEM (GIBCO Cat# 11090-081) supplemented with 10% FBS (Atlas Biologicals Cat# FP-0500-A), 30mM glucose, 1x penicillin-streptomycin solution (GIBCO Cat# 15070-063). P0 mouse or rat hippocampi were dissected in ice cold Hank’s Balanced Salt Solution (HBSS; Cat# GIBCO 14185-052) supplemented with 2.5mM HEPES pH7.4, 30mM glucose, 1mM CaCl_2_, 1mM MgSO_4_ and 4mM NaHCO_3_. Hippocampi were dissociated by incubating in papain solution (20 units/ml papain (Worthington Cat# 3126), 82mM Na_2_SO_4_, 30mM K_2_SO_4_, 5.8mM MgCl_2_, 0.25mM CaCl_2_, 1mM HEPES pH 7.4, 20mM glucose and 0.2mM NaOH) for 30 minutes at 37°C followed by trituration through a polished glass pipette. Dissociated cells were rinsed and resuspended in neuron plating media: MEM supplemented with 10% horse serum (LifeTechnologies Cat# 26050-088), 20mM glucose, 1x penicillin-streptomycin solution, 2mM Glutamax (GIBCO Cat#35050-061), and 1mM sodium pyruvate. Dissociated hippocampal cells were electroporated with 1 pulse at 340V for 900μs pulse length using a square wave protocol in a cuvette with a 2mm electrode (BTX, ECM830, Harvard Apparatus Cat# 45-0002) using the appropriate plasmids and plated at 1×10^5^ cells/well onto glia coated coverslips in a standard 12-well plate. Neurons were kept in a 37°C incubator maintained with 5% CO_2_. A few hours after plating, the neuron plating media was replaced with hippocampal feeding media: Neurobasal A (GIBCO Cat# 10888-022) supplemented with 1x B-27 (GIBCO Cat# 17504-044), 0.2x penicillin-streptomycin solution, 0.5mM Glutamax, 12mM glucose and 1μM AraC (Sigma-Aldrich Cat# C6645).

### Gain-of-function synapse induction assay

Neurons were transfected with GFP-pBOS or co-transfected with GFP-pBOS + FLAG-Kirrel3-pBOS constructs. Equal amounts of DNA were transfected for each condition. Cultures were harvested at 14 days in vitro (DIV) and stained with mouse-anti-GFP, rabbit-anti-Foxp1, guinea pig-anti-VGluT1 and goat-anti-synaptoporin. Transfected CA1 neurons, identified by GFP expression, strong expression of the marker Foxp1 and pyramidal morphology, were imaged at 63x magnification on a Zeiss LSM-7 confocal microscope. Maximum intensity projected images were analyzed in ImageJ. The number of VGluT1 only (originating predominantly from principle excitatory neurons of the CA3, CA2 and CA1 fields combined), SPO only (originating from interneurons) and VGluT1 + SPO co-labelled puncta (originating from DG neurons) were counted along CA1 dendrites and the density was calculated as the number of puncta per length of dendrite. Analysis was performed blind to condition.

### Neuron/HEK293 co-culture assays

Neuron-HEK293 co-cultures were prepared according to an established method (Biederer and Scheiffele, 2007). Briefly, P0 rat hippocampal neurons were plated to poly-D-lysine (PDL; 10μg/cm^2^; Millipore Cat# A-003-E) coated coverslips at 5×10^4^ neurons per well in a 24-well plate. HEK293T cells (ATCC Cat# CRL-3216; RRID: CVCL_0063) were transfected with either GFP-pBOS, GFP-pBOS + FLAG-Neuroligin-1 pCDNA, GFP-pBOS + FLAG-Neurexin-1β-AS4(-)-pCDNA or GFP-pBOS + FLAG-Kirrel3-pBOS (wild-type) 24 hours before seeding to neurons. In one condition for the presynaptic assay, the neurons were electroporated with FLAG-Kirrel3-pBOS before plating.

Test for pre-synapse induction: Transfected HEK293 cells were seeded to 6 DIV neurons at 3×10^4^ cells per well. 24 hours later co-cultures were fixed and immunostained with mouse-anti-FLAG, guinea pig-anti-VGluT1 and goat-anti-GFP. Transfected HEK293 cells in proximity to VGluT1 positive axons were imaged and analyzed. To determine the presynaptic hemisynapse density, the total area of VGluT1 puncta on or within 0.5μm of the edge of each HEK293 cell was quantified and normalized to the total area of the cell in ImageJ. Analysis was performed blind to condition.

Test for post-synapse induction: Transfected HEK293 cells were seeded to 14 DIV neurons at 3×10^4^ cells per well. 24 hours later co-cultures were fixed and immunostained with mouse-anti-FLAG, guinea pig-anti-VGluT1, mouse-anti-PSD-95 and goat-anti-GFP. HEK293 cells in proximity to PSD-95 positive dendrites were imaged and analyzed. To determine the postsynaptic hemisynapse density, the total area of PSD-95 positive, VGluT1 negative puncta on or within 0.5μm of the edge of each HEK293 was quantified and normalized to the total area of the cell in ImageJ. Analysis was performed blind to condition.

### Axon contact assay

P0 Kirrel3 knockout neurons were dissociated and transfected for two conditions. For condition 1 (Kirrel3 cells plus control cells): 2.5×10^4^ neurons per well were co-transfected with GFP-pBOS + FLAG-Kirrel3-pBOS and another 2.5×10^4^ neurons per well were transfected with mCherry-pBOS. For condition 2 (Kirrel3 cells plus Kirrel3 cells): 2.5×10^4^ of the neurons per well were co-transfected with GFP-pBOS + FLAG-Kirrel3-pBOS and 2.5×10^4^ neurons per well were co-transfected with mCherry-pBOS + FLAG-Kirrel3-pBOS. In both conditions, transfected neurons were plated with 5×10^4^ untransfected neurons per well. Neurons were fixed at 7 DIV. GFP-positive dendrites contacting mCherry-positive axons were imaged at 63x and analyzed in ImageJ. To compare the relative amount of mCherry positive axon contact with GFP positive neurons between the conditions, we defined an axon contact index. We created a GFP-positive cell mask by thresholding the GFP channel and an axon mask by thresholding the mCherry channel. The area of mCherry axon mask within in the GFP cell mask was measured. Analysis was performed blind to condition.

### Synapse localization assay

P0 rat hippocampi were dissociated and electroporated with wild-type mCherry-2A-FLAG-Kirrel3-pBOS, or the missense variants, and cultured as described above. At 21 DIV, neurons were live-labelled with chicken-anti-FLAG, fixed, and immunostained with rabbit-anti-2A, guinea pig-anti-VGluT1 and mouse-anti-PSD-95. mCherry-expressing neurons were imaged at 63x and analyzed in ImageJ. Synapses were determined by the juxtaposition of VGluT1 and PSD-95 puncta. The number of synapses was quantified along the dendrites and scored as either FLAG-Kirrel3 positive or FLAG-Kirrel3 negative to determine the percent of synapses with FLAG-Kirrel3. Analysis was performed blind to condition.

### Surface expression assay

Chinese hamster ovarian-K1 (CHO) cells (ATCC, Cat# CCL-61, RRID: CVCL_0214) were plated on PDL-coated (10μg/cm^2^), 18mm coverslips in a 12-well plate at 1×10^5^ cells per well. 24 hours later, cells were transfected with GFP-pBOS or GFP-pBOS + FLAG-Kirrel3-pBOS constructs. Another 24 hours later, cells were live-labelled with chicken-anti-FLAG before permeabilization to detect surface expressed FLAG-Kirrel3. Cells were then fixed, permeabilized and stained with mouse-anti-FLAG to detect intracellular Kirrel3 along with goat-anti-GFP to label transfected cells. The integrated density of the chicken-anti-FLAG (extracellular) signal was quantified per cell and divided by the integrated density of the mouse-anti-FLAG (intracellular) signal to determine the extracellular/intracellular ratio. General culture conditions and solutions for CHO cells were previously described (Martin et al., 2015; Basu et al., 2017). Analysis was performed blind to condition.

### Cell aggregation assay

The aggregation assay was performed as previously described (Martin et al., 2015; Basu et al., 2017). CHO cells were transfected with mCherry-pBOS, wild-type mCherry-2A-FLAG-Kirrel3-pBOS, or the missense equivalents. The aggregation index was calculated by dividing the mCherry integrated density in aggregated cell clumps by the total mCherry integrated density in the well. Analysis was performed blind to condition.

### Immunoblotting

Immunoblotting was conducted by standard techniques. Briefly, CHO cells or HEK293 cells were transfected as needed in a 12-well plate. Cells were collected in 400μL of Laemmli buffer, boiled, run on a 10% Tris-glycine gel, and transferred to a nitrocellulose membrane. The membrane was probed with mouse-anti-Kirrel3 (Neuromab, Cat# 75-333, RRID: AB_2315857) and mouse-anti-GAPDH (Millipore, Cat# AB2302, RRID: AB_ 10615768). Secondary antibody was goat-anti-mouse-HRP (Jackson ImmunoResearch, Cat# 115-035-003, RRID: AB_10015289).

### Experimental Design and Statistical Analysis

Sample sizes were estimated a priori based on results from previous studies (Williams et al, 2011; Martin et al., 2015; Basu et al., 2017). Statistics were computed using Graphpad Prism (RRID: SCR_002798). All data were tested for normality using a D’Agostino and Pearson omnibus normality test and then appropriate parametric or non-parametric tests were used and are reported in the results. For all multiple comparisons, we report the multiplicity adjusted p-values. Graphs show mean +/− SEM. A p-value < 0.05 was considered significant for all tests. *p<0.05, **p<0.01 ***p<0.001, ****p<0.0001. Data from separate experimental trials were normalized to the negative control for the experimental trial before combining datasets to control for culture to culture variability. For k-means clustering we used the mean rank values for each condition from the synapse induction assay and the cell aggregation assay consistent with the non-parametric tests used to compare them. Mean ranks were normalized to the wild-type mean rank for ease of viewing.

## RESULTS

### Exogenous Kirrel3 expression induces ectopic synapse formation between incorrect targets

Kirrel3 knockout results in selective synapse loss but it is unknown if Kirrel3 plays an instructive role in selective synapse development. To complement our previous loss-of-function studies, we sought to test if exogenous Kirrel3 expression induces synapse formation via a gain-of-function assay. To this end, we developed an in vitro assay based on the fact that target specific synapse formation is largely maintained between cultured hippocampal neurons (Williams et al., 2011). Previously it was shown that cultured DG neurons frequently synapse with CA3 neurons (an appropriate target in vivo) but only rarely synapse with CA1 neurons (an inappropriate target in vivo) despite the fact that DG axons similarly contact CA3 and CA1 dendrites in culture (Williams et al., 2011). Because CA1 neurons do not appreciably express Kirrel3, we used this in vitro specificity to our advantage and tested if exogenous Kirrel3 expression in CA1 neurons could induce ectopic DG-to-CA1 synapses (Fig. 1B).

To accomplish our goal, we needed to unambiguously identify CA1 neurons and DG synapses in culture. We demonstrate that CA1 neurons are identified by high expression of the transcription factor Foxp1, while pre-synapses arising from DG axons are identified by co-expression of the presynaptic proteins synaptoporin (SPO) and vesicular glutamate transporter 1 (VGluT1) (Fig. 1C; Ferland et al., 2003; Williams et al., 2011;). This combination of markers allows us to quantify the number of DG pre-synapses formed onto CA1 dendrites. We transfected dissociated rat hippocampal neurons with GFP alone or GFP plus FLAG-Kirrel3 and quantified the number of DG pre-synapses per unit length of CA1 dendrite to determine the DG pre-synapse density in GFP-positive neurons (Fig. 1D-E). Normally, cultured CA1 neurons have few DG pre-synapses but exogenous expression of Kirrel3 in CA1 neurons significantly increases the density of DG pre-synapses onto CA1 neurons by more than two-fold (Fig. 1E; Mann-Whitney U=570, p=0.0027). This effect is specific for DG synapses as we do not observe an increase in the number of synapses arising from other types of glutamatergic neurons in the culture. This includes CA3, CA2, and CA1 neurons, which are collectively marked by expression of VGluT1, but not synaptoporin, and are referred to here as CA synapses (Fig. 1F; Mann-Whitney U=905, p=0.9552).

Kirrel3 mediates homophilic, trans-cellular adhesion in vitro, and in vivo loss of function studies suggest that Kirrel3 mediates synapse formation selectively between Kirrel3-expressing neurons (Gerke et al, 2005; Serizawa et al, 2006; Martin et al., 2015). Together, this suggests Kirrel3’s function requires trans-cellular interactions with other Kirrel3 molecules. Since all DG neurons normally express Kirrel3, we next tested if Kirrel3’s gain-of-function effect depends upon the presence of endogenous Kirrel3 in DG axons. To do this, we performed the same assay in hippocampal neurons cultured from both Kirrel3 wild-type and knockout mice, taking extra precaution to not quantify CA1 neurons that were contacted by transfected axons (marked by GFP). Similar to our results in rat neurons, we find that exogenous Kirrel3 expression potently induces ectopic DG-to-CA1 synapses in cultures prepared from wild-type mice, but Kirrel3 has no synaptogenic activity in cultures prepared from Kirrel3 knockout mice (Fig. 1G; two-way ANOVA, F_(1,111)_=5.049, interaction, p=0.0266; Sidak’s multiple comparison WT-GFP vs WT-Kirrel3 p=0.0088, KO-GFP vs KO-Kirrel3 p=0.9560). This strongly suggests that in our gain-of-function assay postsynaptic Kirrel3 in the CA1 dendrite interacts with pre-synaptic Kirrel3 in the DG axons to induce synapse formation via homophilic, trans-cellular interactions (Fig. 1H).

### Kirrel3-induced synapses do not result from increased axon-dendrite contact

It is possible Kirrel3 interactions directly instruct neurons to recruit synaptic components at existing axon-dendrite contact points. Alternatively, Kirrel3 expression may increase the amount of axon-dendrite contact between DG and CA1 neurons, and thereby increase the chance that synapses will form via other molecules. Thus, we next tested if Kirrel3 expression increases axon-dendrite contacts. Tools to identify endogenously-expressing Kirrel3 positive and negative axons in culture are not available, so we cultured neurons from Kirrel3 knockout mice and transfected Kirrel3 in a controlled manner. Knockout neurons were co-transfected with GFP and Kirrel3 and mixed with knockout neurons transfected with either mCherry, or mCherry and Kirrel3. Thus, while in both conditions GFP-labelled neurons expressed Kirrel3: in one condition they were contacted by mCherry axons that were Kirrel3 negative, and in the other condition they were contacted by mCherry axons that were Kirrel3 positive. Quantification of the overlap between GFP dendrites and mCherry axons revealed a similar amount of axon-dendrite overlap regardless of whether the axons expressed Kirrel3 or not (Fig. 2A-2B; Mann-Whitney U=257, p=0.2445). This suggests Kirrel3 expression does not induce synapses simply by increasing axon-dendrite contact and supports a model in which Kirrel3 trans-cellular interactions actively recruit synaptic factors to existing points of axon-dendrite contacts.

**Figure 2.**
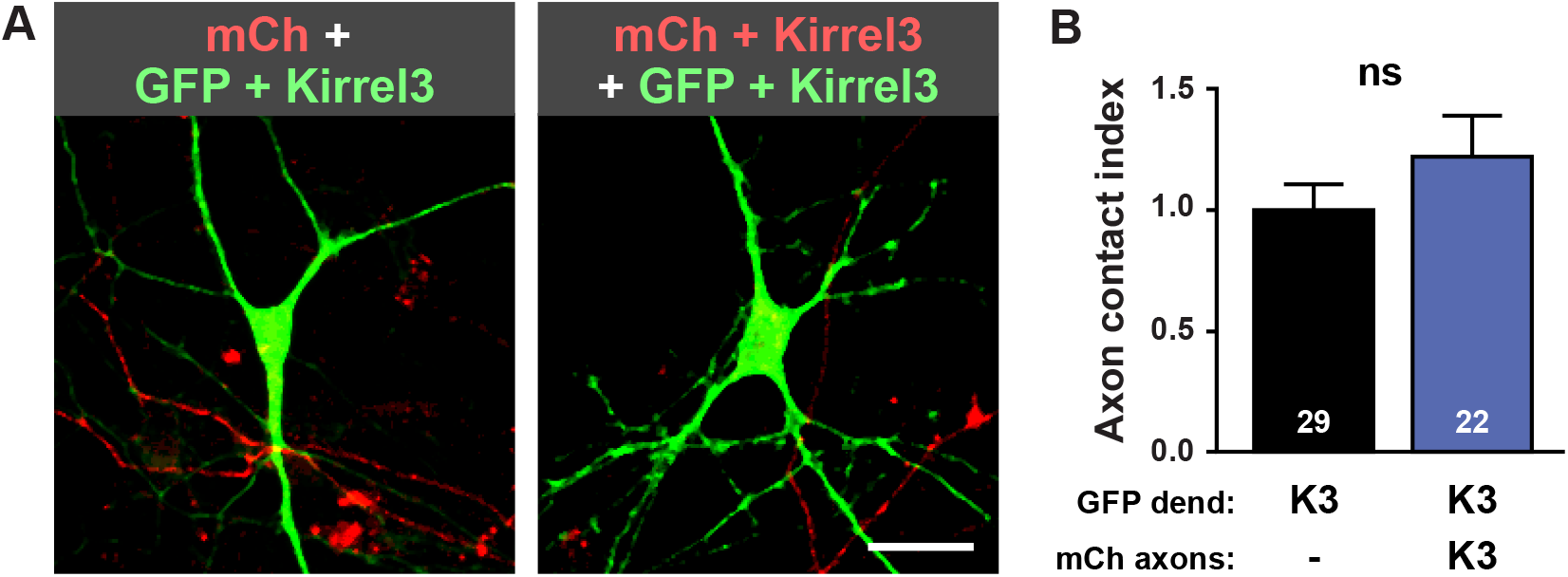
Kirrel3 expression does not increase axon-dendrite contact in cultured neurons. **A,**Representative images of neurons transfected with GFP and Kirrel3 (green) contacted by mCherry (mCh) labeled axons with (top) or without (bottom) Kirrel3. Scale bar = 10μm. **B**, Quantification of axon contact assay normalized to the Kirrel3-negative mCherry-labelled axon condition. Axon contact index is the area of mCherry positive axon per area GFP positive cell. n=22-29 neurons (indicated in each bar) from 2 cultures, p=0.2445 (Mann Whitney Test). ns = not significant.

### Kirrel3-mediated synaptogenesis requires post-synaptic neurons

We next tested if Kirrel3 is sufficient to induce synapses using a well-established co-culture assay in which synaptogenic proteins are presented to neurons in non-neuronal cells (Scheiffele et al. 2000, Graf et al., 2004; Biederer and Scheiffele, 2007). In this assay, neurons abundantly contact HEK293 cells but do not recognize HEK293 cells as synaptic partners. However, if HEK293 cells express some postsynaptic molecules such as Neuroligin-1, the neurons form presynapse structures, identified by VGlut1, on the HEK293 cells (Fig. 3A) (Scheiffele et al., 2000). Conversely, when HEK293 cells express some presynaptic molecules such as Neurexin-1, the neurons form postsynapse structures, identified by PSD-95, on the HEK293 cells (Graf et al., 2004). Interestingly, we find that when HEK293 cells express Kirrel3, the neurons form neither pre-nor post-synapses with HEK293 cells, though both of our positive controls (Neurexin-1 and Neuroligin-1) functioned as expected (Fig. 3A-3D; Pre-synapses: Kruskal-Wallis test with Dunn’s multiple comparisons GFP vs Kirrel3 p=0.3988, GFP vs Neuroligin-1 p=0.0074; Post-synapses: Kruskal-Wallis test with Dunn’s multiple comparisons GFP vs Kirrel3 p>0.9999, GFP vs Neurexin-1 p=0.0252). Because Kirrel3 expression specifically induces DG synapses, we also conducted this assay using the DG-specific marker synaptoporin instead of the generic marker VGluT1. Unfortunately, we find synaptoporin is not reliably expressed in the 7 DIV neurons used for this assay. We circumvented this limitation by transfecting extra Kirrel3 into hippocampal neurons and then quantifying the density of presynaptic puncta on Kirrel3-expressing HEK293 cells contacting Kirrel3-transfected axons. We found that this still had no effect on the density of pre-synapses formed onto HEK293 cells (Fig. 3A-B; Kruskal-Wallis test with Dunn’s multiple comparisons GFP vs Kirrel3 neuron and HEK293 expression p>0.9999). In conclusion, we find Kirrel3 is not sufficient to induce synapses when expressed in a non-neuronal cell. Taken together, our work indicates that Kirrel3’s synaptogenic function requires both trans-cellular Kirrel3 binding and interactions with an additional co-factor(s) present in CA1 neurons but not in HEK293 cells.

**Figure 3.**
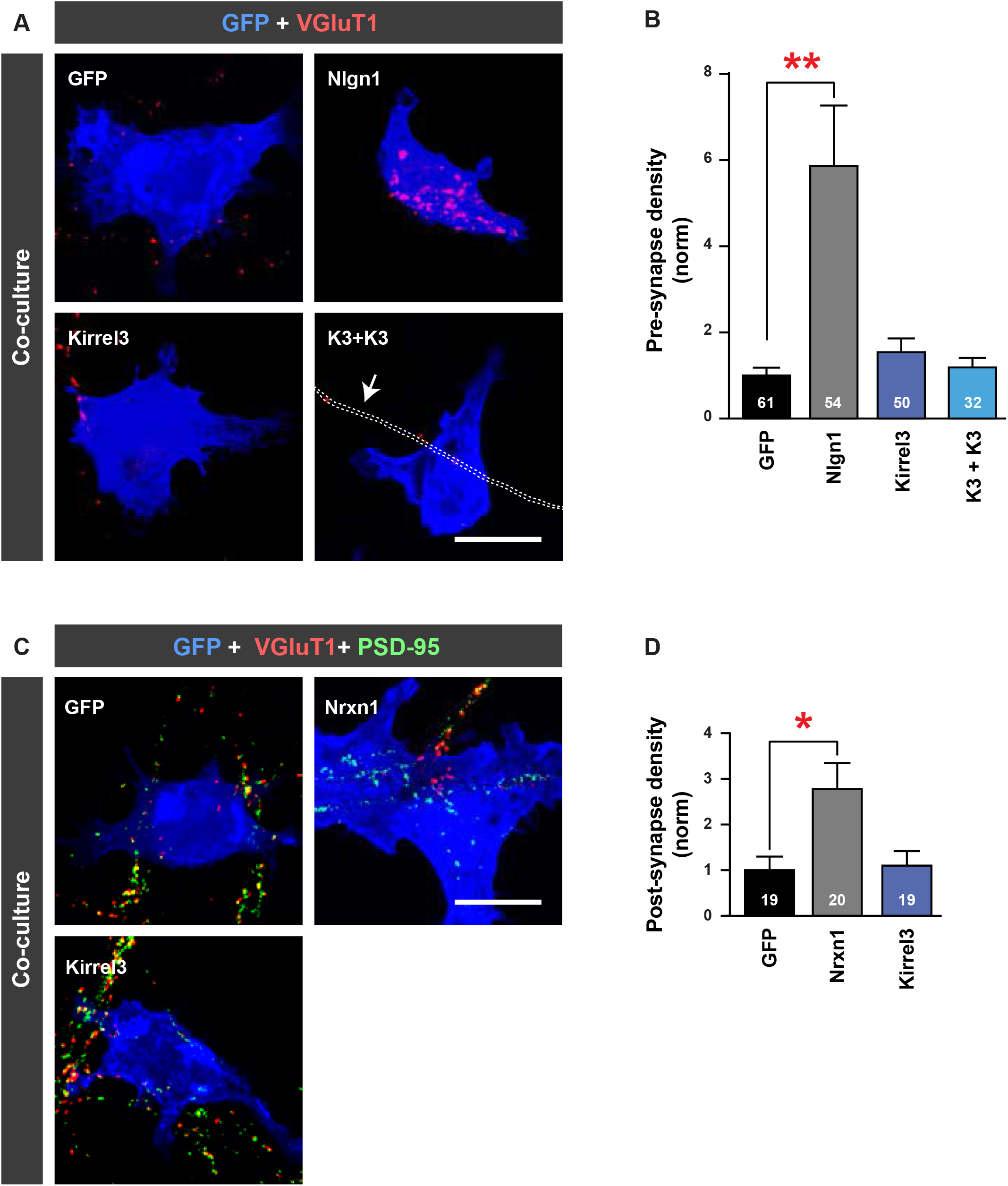
Kirrel3 in a non-neuronal cell is not sufficient to mediate synapse formation. **A**, Representative images of the presynaptic neuron/HEK293 co-culture assay. Cultures were immunostained for GFP (blue) to label transfected HEK293 cells and VGluT1 (red) to label pre-synapses. Transfection conditions are indicated on each image. Nlgn1 = Neuroligin-1, K3 = Kirrel3. Arrow in bottom right panel points to an outline of a FLAG-Kirrel3 positive axon **B**, Quantification of presynaptic hemisynapse density from the co-culture assay normalized to the GFP-only negative control (GFP). n=32-61 cells (indicated in each bar) from 2 cultures, GFP vs Nlgn1 p=0.0074; GFP vs Kirrel3, p=0.3988; GFP vs K3+K3 p>0.9999 (One-way ANOVA Kruskal-Wallis test, p=0.0226, with Dunn’s multiple comparisons). **C,** Images of the postsynaptic hemisynapse density from the co-culture assay. Cultures were immunostained with GFP (blue), vGluT1 (red) and PSD-95 (green). Postsynaptic hemisynapses are labelled by PSD-95 alone with no VGluT1. Nrxn1 = Neurexin-1. **D**, Quantification of postsynaptic hemisynapse density from co-culture assay normalized to the GFP-only negative control. n=19-20 cells (indicated in each bar) from one culture, GFP vs Nrxn1 p=0.0252; GFP vs Kirrel3 p>0.9999 (One-way ANOVA Kruskal-Wallis test, p=0.0256, with Dunn’s multiple comparisons). Scale bar in A and C = 20μm.

### Kirrel3 missense variants appropriately traffic to synapses and the cell surface

Thus far, we used our in vitro gain-of-function assay to identify basic mechanistic principles of Kirrel3 signaling during synapse formation. Next, we tested if and how disease-associated Kirrel3 missense variants affect its function. Accordingly, we cloned six missense variants identified as possible risk factors for neurodevelopmental disorders (Fig. 4A; Table 1; Bhalla, et al., 2008; De Rubeis, et al., 2014; Iossifov et al., 2014; Yuen, et al., 2016). These variants span the length of the Kirrel3 protein and are fully conserved between the mouse and human proteins, which share 98% amino acid identity (Gerke et al., 2005). Most variants are found in patients with both autism and intellectual disability. One variant, M673I, has not been reported previously. We identified this as a likely pathogenic variant in a male patient with developmental delay, intellectual disability, ADHD, and obesity (see methods for complete description).

**Figure 4.**
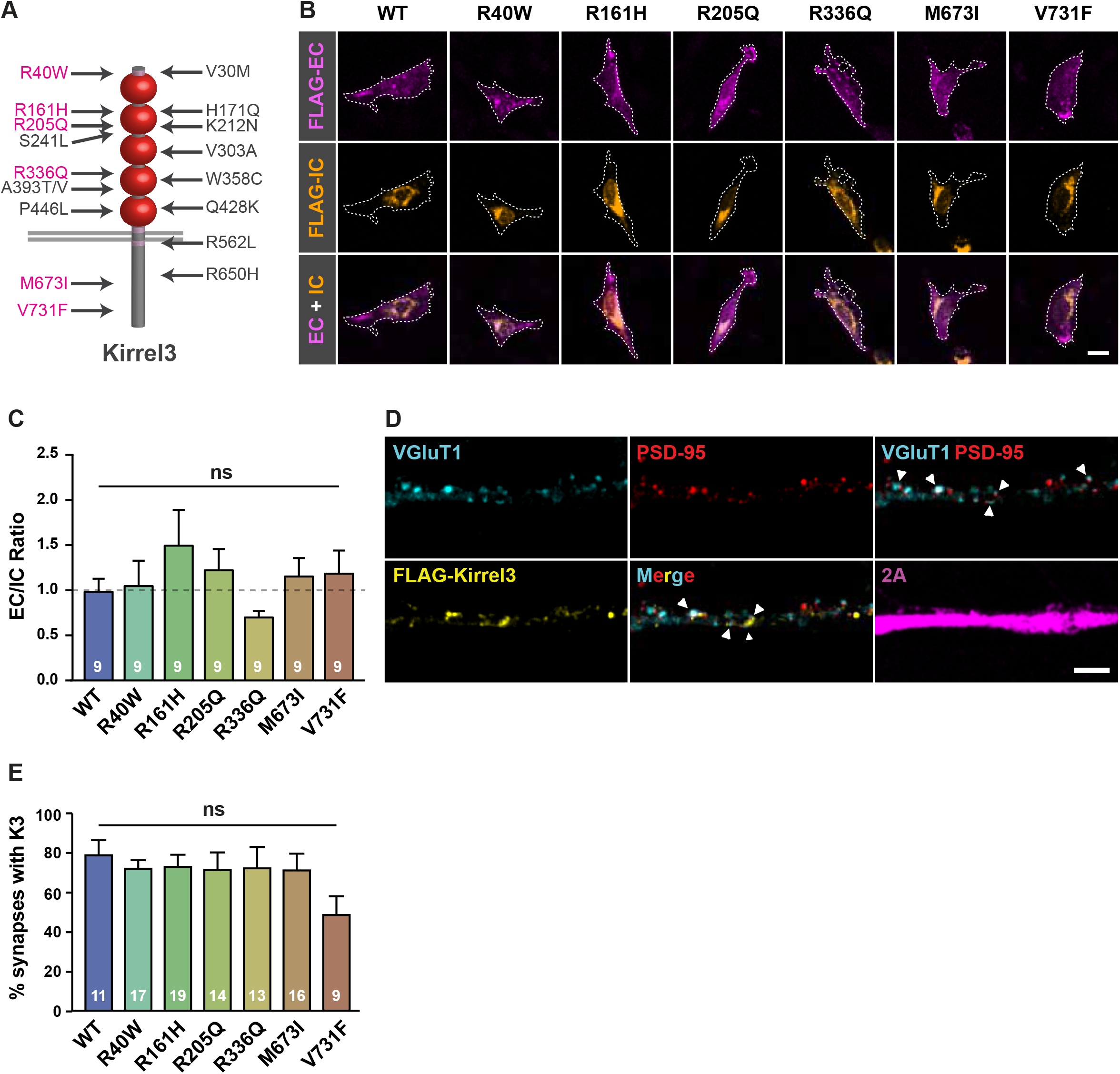
Kirrel3 missense variants show normal surface localization. **A**, Schematic of Kirrel3 protein with approximate positions of disease-associated missense variants. Variants tested in this study are magenta. **B**, Representative images of CHO cells transfected with mCh-2A-FLAG-Kirrel3 variants. Cells were live-labelled with chicken-anti-FLAG (magenta) to show extracellular (EC) Kirrel3 protein and then permeabilized and stained with mouse-anti-FLAG (orange) to show intracellular (IC) Kirrel3 protein. Cell area was determined by GFP expression and is depicted by a white outline. Scale bar = 15μm. **C**, Quantification of EC/IC Kirrel3 protein levels normalized to wild-type Kirrel3. n=9 cells each from 3 cultures, p=0.4424 (One-way ANOVA). **D**, Representative images showing a dendrite from a mCherry-2A-FLAG-Kirrel3 wild-type transfected neuron. EC FLAG-Kirrel3 (yellow) localizes adjacent to juxtaposed synaptic markers VGluT1 (blue) and PSD-95 (red). mCherry-2A (magenta) serves as a cell marker. Scale bar = 4μm. **E**, Quantification of the percent of synapses (juxtaposed VGluT1^+^, PSD-95^+^ puncta) with adjacent FLAG-Kirrel3 proteins for each variant. n=9-19 neurons (indicated in each bar) from two cultures, p=0.3197 (One-way ANOVA Kruskal-Wallis test). ns = not significant

Missense variants can impair the function of synaptic molecules by altering their trafficking or surface expression (Chih et al., 2004; Zhang et al., 2009; Jaco et al., 2010). Thus, we first tested if Kirrel3 variants are expressed on the cell surface similar to wild-type Kirrel3. We transfected CHO cells with Kirrel3 plasmids containing an extracellular FLAG tag that does not interfere with Kirrel3 homophilic binding (Martin et al., 2015). The cells were live-labelled with chicken-anti-FLAG antibodies to label Kirrel3 protein on the extracellular surface. Then, cells were fixed, permeabilized, and incubated with mouse-anti-FLAG antibodies to label intracellular Kirrel3 (Fig. 4B). The ratio of extracellular to intracellular Kirrel3 was quantified for individual cells. We find the relative intensities of extracellular to intracellular FLAG-Kirrel3 are indistinguishable between groups, indicating that all six missense variants are appropriately trafficked to the cell surface similar to wild-type Kirrel3 (Fig. 4C; one-way ANOVA F_(6,56)_=0.9878, p=0.4424).

Wild-type Kirrel3 localizes at or near excitatory synapses (Martin et al., 2015; Roh et al, 2017). Thus, we tested if Kirrel3 missense variants also localize to the cell surface at synapses in neurons. We expressed mCherry-2A-FLAG Kirrel3 and missense variants in cultured neurons and determined their localization relative to synapses by co-staining for surface-expressed FLAG, VGluT1, and PSD-95 (Fig. 4D). All Kirrel3 variants display a punctate expression pattern and are found in close proximity to juxtaposed VGluT1/PSD-95 puncta at similar rates (Fig. 4E; Kruskal-Wallis test =7.013; p=0.3197). Collectively, we do not observe general trafficking defects in disease-associated Kirrel3 missense variants.

### Disease-associated Kirrel3 variants attenuate Kirrel3 function

We next tested the ability of each variant to induce DG pre-synapses when ectopically expressed in CA1 neurons compared to wild-type (as shown Figure 1D). Importantly, we found that four of the six disease-associated variants (R161H, R205Q, R336Q and V731F) significantly attenuate Kirrel3 mediated pre-synapse formation when compared to wild-type Kirrel3 (Fig. 5A-B, Kruskal-Wallis test with Dunn’s multiple comparisons WT vs GFP p=0.<0.0001, WT vs R40W p=0.2495, WT vs R161H p=0.0472, WT vs R205Q p=0.0218, WT vs R336Q p=0.0135, WT vs M673I p=0.9844, WT vs V731F p=0.0032). Another variant, R40W, appears to attenuate synapse formation but does not achieve statistical significance in this comparison. We then applied statistical tests to determine the ability of the missense variants to induce pre-synapse formation relative to negative control neurons transfected with GFP alone using the same dataset. Consistent with our previous analysis, DG-pre-synapse densities on CA1s expressing R161H, R205Q, R336Q or V731F are not significantly different from GFP-only transfected neurons, suggesting that these variants do not induce synapse formation (Fig. 5C; Kruskal-Wallis test with Dunn’s multiple comparisons GFP vs WT p<0.0001, GFP vs R161H p>0.9999, GFP vs R205Q p>0.9999, GFP vs R336Q p>0.9999, GFP vs M673I p=0.0382, GFP vs V731F p>0.9999). In addition, we found that the R40W variant is also not significantly different from GFP transfected controls (Kruskal-Wallis test with Dunn’s multiple comparisons GFP vs R40W p=0.1299), suggesting that this variant also does not induce synapse formation. The different outcomes for this variant depending on the statistics applied suggest that the R40W variant likely attenuates Kirrel3 function very near the sensitivity threshold of this functional assay. Thus, we conclude that five of the six tested disease-associated Kirrel3 variants have impaired synaptogenic function. Our results provide the first functional evidence that Kirrel3 missense variants may cause disease through the selective loss of synapses.

**Figure 5.**
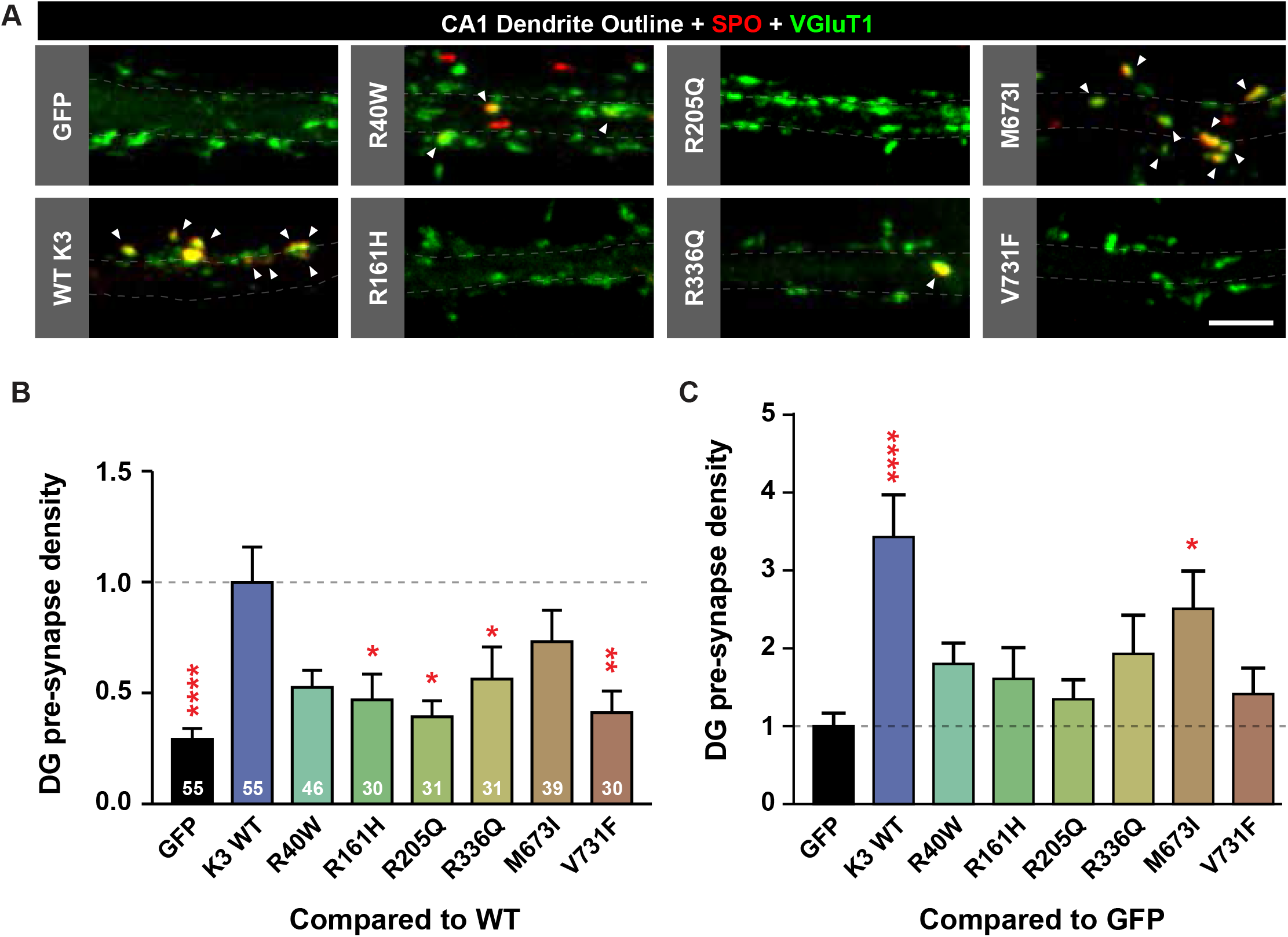
Missense variants attenuate Kirrel3-mediated synapse formation. **A**, Representative images of CA1 dendrites transfected with GFP only (GFP), or GFP and indicated FLAG-Kirrel3 variants. Outlines show dendrite area as determined from GFP expression. DG pre-synapses (yellow) are identified by co-labelling of SPO (red) and VGluT1 (green). Arrows point to DG pre-synapses. Scale bar = 5μm. **B**, Quantification of DG pre-synapse density on CA1 dendrites normalized to wild-type Kirrel3 and with multiple comparisons to wild-type Kirrel3. n=30-55 neurons (indicated in each bar) from 3-5 cultures, (One-way ANOVA Kruskal-Wallis test with multiple comparisons to WT Kirrel3). **C**, Same data as ssssspresented in B but shown normalized to GFP and with multiple comparisons to GFP. See results section for specific p-values but *p<0.05, **p<0.01, and ****p<0.0001.

### Kirrel3 missense variants fail to induce pre-synapses via different mechanisms

Missense variants affecting Kirrel3 function span the extracellular and intracellular domains suggesting they may impair Kirrel3 function in distinct ways (Fig. 4A, Table 1). We reasoned that variants in the extracellular domain may eliminate Kirrel3 homophilic binding while variants in the intracellular domain may alter downstream signaling. It is not yet possible to test Kirrel3 intracellular signaling because very little is known about Kirrel3’s intracellular interactions or signaling mechanisms. Therefore, we tested the ability of each missense variant to mediate homophilic binding in trans using an established CHO cell aggregation assay. CHO cells do not endogenously express appreciable levels of Kirrel3 (Fig. 6A) and when mixed in suspension they do not normally form cell aggregates (Fig. 6B-C). In contrast, CHO cells transfected with wild-type Kirrel3 form multicellular aggregates consistent with Kirrel3 undergoing homophilic, trans-cellular binding (Fig. 6B-C; Martin et al., 2015). After testing each variant, we find that cells transfected with R40W, R161H, R336Q and M673I aggregate similarly to wild-type Kirrel3, but the R205Q and V731F missense variants do not induce cell aggregation (Fig. 6C, Kruskal-Wallis test with Dunn’s multiple comparisons WT vs Ctrl p<0.0001, WT vs R40W p>0.9999, WT vs R161H p>0.9999, WT vs R205Q p<0.0001, WT vs R336Q p>0.9999, WT vs M673I p>0.9999, WT vs V731F p=0.0042). Subsequent k-means clustering defines four groups of variants with respect to synaptogenic and aggregation ability (Fig. 6D). Wild-type Kirrel3 and the M673I variant group together as they exhibit homophilic trans-cellular binding and induce synapse formation to a greater extent than the other variants. The R40W variant is isolated reflecting its intermediate ability to induce synapse formation with normal trans-cellular binding. R161H and R336Q form a group characterized by attenuated synaptogenic ability with normal trans-cellular binding. Finally, R205Q and V731F group with the negative control as they are completely binding deficient and do not induce synapse formation. Collectively these results indicate that missense variants impact Kirrel3 function in distinct ways: some attenuate Kirrel3 mediated trans-cellular interactions, while others maintain an ability to mediate contact recognition but still fail to induce synaptogenesis strongly implicating additional unidentified signaling mechanisms.

**Figure 6.**
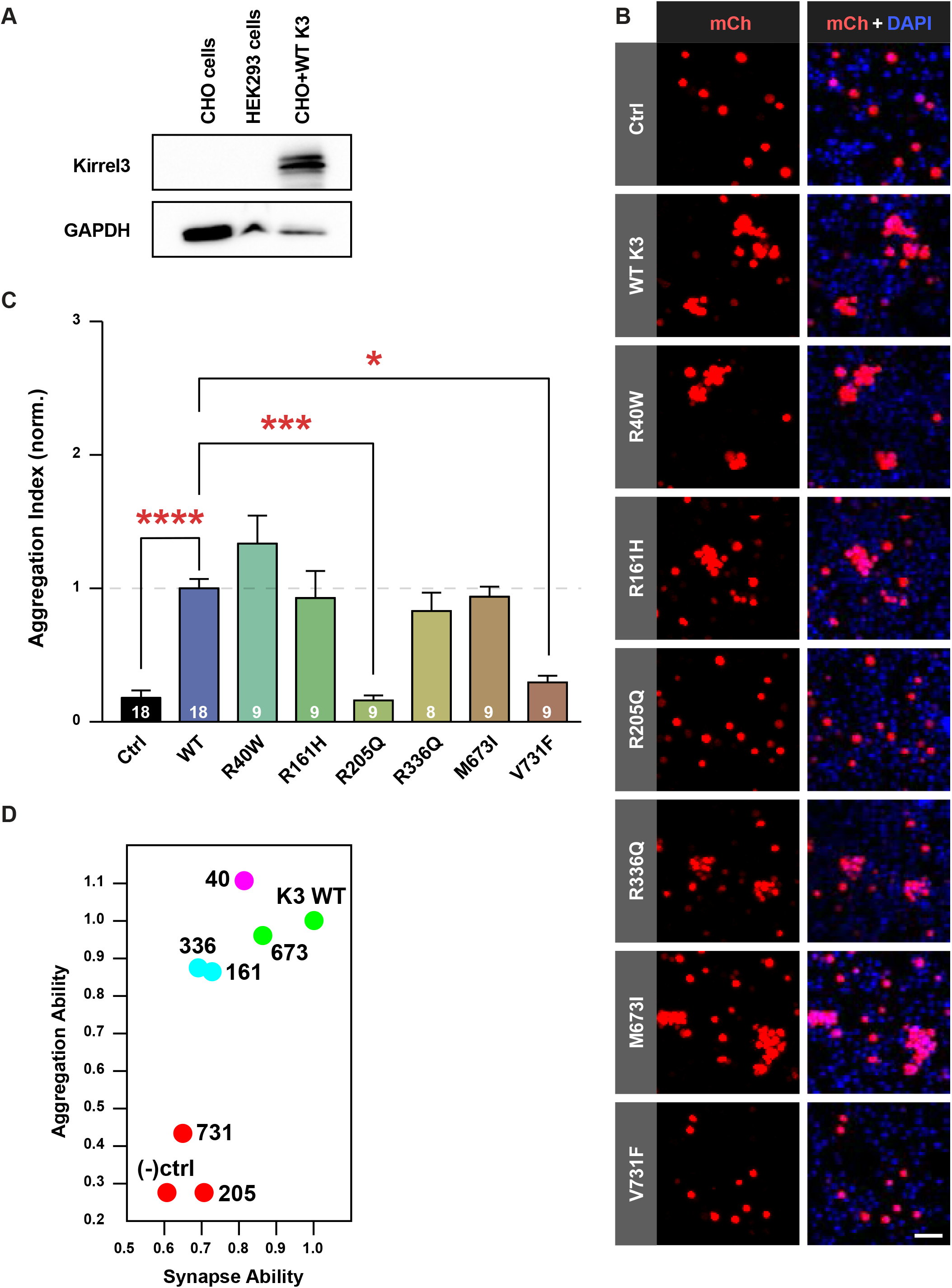
Kirrel3 missense variants fail to induce synapse development via different mechanisms. **A**, Immunoblot of untransfected CHO and HEK293 cell lysates showing no detectable endogenous Kirrel3 expression. CHO cells transfected with FLAG-Kirrel3-pBOS were used as a positive control. **B**, Representative images from CHO aggregation assay showing CHO cells transfected with mCherry alone as a negative control (Ctrl), mCherry-2A-Kirrel3-wild-type (WT K3), or mCherry-2A-Kirrel3 variants. mCherry (mCh) signal is shown in red, DAPI in blue. Scale bar = 50μm. **C**, Quantification of CHO cell aggregation assay normalized to wild-type Kirrel3. n=8-18 wells (indicated in each bar) from 3 cultures. See results section for specific p-values. *p<0.05, ***p<0.001, ****p<0.0001. **D**, K-means clustering of wild-type Kirrel3, missense variants, and negative control based on their ability to aggregate CHO cells and induce DG pre-synapse formation using mean rank values normalized to wild-type Kirrel3 for each condition from the two assays presented in figures 5B and 6C.

## DISCUSSION

### Kirrel3 functions in synapse development as a homophilic cell-recognition tag

Most synapses do not form randomly between any neurons that come into contact with each other. Rather, neurites coursing through a dense microenvironment often select specific partners among a milieu of incorrect choices. A long-standing hypothesis proposes that neuronal wiring requires matching affinity tags selectively expressed on pre- and postsynaptic neurons (Sperry, 1963; Benson et al., 2001; Sanes and Yamagata, 2009; Südholf, 2018). The identity of these tags and how they signal at specific synapses remain largely unresolved.

Here, we directly tested if Kirrel3 acts as synaptogenic, cell-recognition tag using a novel gain-of-function assay. We demonstrate that ectopic Kirrel3 expression in CA1 neurons specifically induces ectopic DG synapse formation, providing the first direct evidence that Kirrel3 plays an instructive role in synapse development. Furthermore, we show that ectopic Kirrel3 has no function when expressed in cultures prepared from Kirrel3 knockout mice. This indicates that ectopically-expressed Kirrel3 function depends on the presence of endogenous Kirrel3, most likely because ectopic Kirrel3 in the CA1 dendrites interacts with endogenous Kirrel3 in DG axons to induce DG synapses. This strongly suggests Kirrel3’s trans-cellular, homophilic interactions are necessary for its function. This is significant because, although prior work indicates that Kirrel3 can undergo homophilic interactions, a functional role of Kirrel3 homophilic interactions at synapses had not been directly tested (Gerk et al., 2005; Serizawa et al., 2006; Martin et al., 2015; Prince et al., 2013; Brignall et al., 2018). Moreover, in the mouse neuromuscular junction, mouse kidney, and at C. elegans synapses, Kirrel3 functions via heterophilic interactions with another Ig-superfamily protein Nephrin-1 (C. elegans Syg-1, Syg-2 respectively) (Shen and Bargmann, 2003; Shen et al., 2004; Gerke et al., 2005; Ding et al., 2007; Chao et al., 2008; Komori et al., 2008; Chia et al., 2014; Özkan et al., 2014). Our study does not rule out the possibility that Kirrel3 binds in trans to other molecules, but we clearly show that Kirrel3-Kirrel3 trans-homophilic interactions are necessary for its synaptogenic function.

### Kirrel3 has multifaceted roles as a recognition, adhesion, and signaling molecule

Cell adhesion proteins are important for contact recognition between neurons but very little is known about downstream signaling events that couple contact recognition to synapse development. Therefore, a common misperception is that adhesive proteins expressed on the cell surface act as a “molecular glue” to hold juxtaposed membranes together long enough for other general synaptogenic factors to be recruited. In contrast, our findings support a model in which Kirrel3 likely mediates a combination of contact recognition, adhesion, and downstream synaptogenic signaling and directly implicate Kirrel3 as a bona fide signaling molecule. Several lines of evidence support this conclusion. First, in support that Kirrel3 likely mediates contact recognition and adhesion, we identified two variants, R205Q and V731F, that cannot bind in trans and have no synaptogenic activity. This supports the conclusion that trans-cellular binding is necessary for Kirrel3 function. Interestingly, the V731F variant resides in the Kirrel3 intracellular domain and we speculate that this variant produces a conformational change that directly inhibits trans-cellular interactions by the extracellular domain. Nonetheless, our results strongly suggest that trans-cellular binding is absolutely critical for Kirrel3 synaptic function. Second, in support of the idea that Kirrel3 has a non-adhesive signaling function, we also identified variants, R161H and R336Q, which bind normally in trans yet have impaired synaptogenic activity. These results indicate that trans-cellular binding is not sufficient for Kirrel3 function and strongly suggest Kirrel3 is involved in synaptic signaling in addition to its role in trans-cellular recognition or adhesion. Additional support for this conclusion comes from the fact that Kirrel3 is not synaptogenic if presented to neurons by a non-neuronal cell. This also strongly suggests that trans-cellular binding is not sufficient for its synaptogenic function and that Kirrel3 function requires a neuronal co-factor. In sum, we find that trans-cellular binding is necessary but not sufficient for Kirrel3 function and that Kirrel3 must interact with other neuronal proteins to induce a synapse. Thus, Kirrel3 does more than act as synaptic glue.

### What proteins might mediate a Kirrel3 signal?

For the presynaptic DG axon to differentiate between a HEK293 cell and neuron, we postulate that Kirrel3 likely interacts with another extracellular or transmembrane molecule in the postsynaptic neuron and that molecule must be lacking in the HEK293 cell. Interestingly, the Kirrel3 extracellular domain was reported to interact with the cell adhesion molecule Neurofascin but this remains to be tested under physiological conditions (Völker et al., 2018).

Kirrel3 intracellular interactions are also likely to be important for Kirrel3 mediated synapse development. Most interestingly, the Kirrel3 intracellular domain ends in a conserved PDZ-binding domain, which is a common protein motif of synaptic organizing molecules. It was suggested that mouse Kirrel3 binds to the PDZ domain proteins CASK, PSD95, and ZO-1 (Huber et al., 2003; Gerke et al., 2006; Roh et al., 2017). Other evidence for putative Kirrel3 binding partners comes from studies using Drosophila and C. elegans. Here, Kirrel3 orthologues are shown to physically or genetically interact with Rols7 (Tanc1/2, a synaptic scaffold), Loner (Brag2, an Arf-GEF), Lin10 (Mint, an active zone component), WVE-1 (WAVE-1, an actin regulator), and SKR-1 (Skp1, an E3 ubiquitin ligase) (Kreisköther et al., 2006; Vishnu et al., 2006; Bulchand et al., 2010; Ding et al., 2007; Chia et al., 2014). However, none of these Kirrel3 interacting proteins have yet been well validated under physiological conditions in the mammalian brain. Further work is needed to identify new Kirrel3 signal transduction candidates and screen existing candidates. The identity of Kirrel3 signaling molecules and whether or not they are shared by other synapse specificity factors will be an important question to address in future studies.

### Kirrel3 variants associated with disease impair synapse formation

Kirrel3 is an emerging hotspot for rare variants linked to neurodevelopmental disorders (Bhalla et al, 2008; Kaminsky et al, 2011; Ben-David and Shifman, 2012; Guerin et al, 2012; Michaelson et al, 2012; Neale et al, 2012; Talkowski et al, 2012; Iossifov et al, 2014; Rubeis et al, 2014; Wang et al, 2016; Yuen et al, 2016; Li et al, 2017; Guo et al, 2019; Leblond, et al, 2019). Here we provide the first functional evidence demonstrating that Kirrel3 variants could cause disease.

Interestingly, not all of the missense variants studied present a phenotype. Our newly reported M673I variant performs similar to wild-type Kirrel3 in all of the assays presented here. While this negative result could mean this particular variant is not disease causing, it is also possible that the variant requires a second mutation in another unidentified gene to impair its function or that our current set of tests for Kirrel3 function have not yet revealed the cellular phenotypes caused by this variant. For example, because CA1 neurons do not normally express Kirrel3, we would not expect this assay to identify dominant negative mutations (discussed further below), only loss-of-function mutations. Thus, M673I could function as a dominant negative and we are in the process of developing new assays to test this class of mutations. Nonetheless, 5 of 6 tested disease-associated Kirrel3 variants have impaired function in our assays, providing the first functional evidence that most disease-associated Kirrel3 variants could cause neurodevelopmental disorders.

Most disease-associated Kirrel3 variants are rare, spontaneous mutations for which patients are heterozygous. If these variants cause disease then Kirrel3 is likely haploinsufficient and/or disease-associated variants act as dominant negative proteins. In support of haploinsufficiency, an interstitial deletion on part of chromosome 11, which leads to a complete loss of one copy of Kirrel3 along with other genes, is associated with Jacobsen syndrome, a developmental disorder with frequent neurological symptoms including ASD, epilepsy, and intellectual disability (Guerin et al., 2012). Furthermore, Kirrel3 has a pLI score, a probability measure that a gene is intolerant to heterozygous loss of function variation, of 0.97 suggesting haploinsufficiency for loss of function variants (gnomAD database https://gnomad.broadinstitute.org/; Lek et al, 2016). On the other hand, we find that all missense variants tested localize to the cell surface at synapses similar to wild-type Kirrel3, suggesting that Kirrel3 variants could interact in a dominant negative fashion by sequestering Kirrel3 binding partners. Future work is needed to determine the precise mechanism of Kirrel3 and each variant in vivo, but results from this study provide a critical foundation that Kirrel3 variants are likely bona fide disease risk factors.

The study of Kirrel3 has strong potential to contribute to our broader understanding of the cellular and circuit defects underlying ASD. Hundreds of genes are implicated as ASD risk factors but relatively few have been the focus of intense study. Many commonly studied ASD risk genes are either ubiquitously expressed throughout the body (TSC1, PTEN, MECP2, CHD8) or by many neurons across most regions in the brain (DSCAM, NLGN3, SYNGAP1, SHANK3). This makes it difficult to identify the essential cell and circuit level changes driving disorders such as ASD and ID, and any future therapeutics based on widely expressed gene pathways are likely to have many adverse side effects. In contrast, Kirrel3 is expressed by a relatively limited and specific subset of cell types (Lein et al., 2007; Allen Institute for Brain Science, 2004; Allen institute for brain science, 2010; Hawrylycz et al., 2012; Martin et al., 2015; Roh et al., 2017). Thus, understanding the mechanism of Kirrel3 in synapse formation and function could point to a core set of synapse and circuit changes underlying some forms of ASD and ID.

## Acknowledgements

This work was supported by grants from the National Institute of Mental Health (R01 MH105426, M.E.W), Brain Research Foundation Fay/Frank Seed Grant (M.E.W), Childcare Foundation (S.E.A), and an Autism Speaks Dennis Weatherstone Predoctoral Fellowship 10116 (E.A.M). We thank the patients and families reported in this study, S. Hawatmeh for technical assistance, the entire Williams lab for experimental help and discussion, and R. Dorsky, J. Christian, and M. Deans for comments on the manuscript.

